# The importance of assigning responsibility during evaluation in order to increase student satisfaction from physical education classes: A model of structural equations

**DOI:** 10.1101/489229

**Authors:** Marta Leyton, Susana Lobato, Marco Batista, Ruth Jiménez

## Abstract

Considering the benefits that students report when evaluating physical education classes, the purpose of the present study was to analyse the relationships between the assignment of student responsibility in the evaluation, motivational variables and the satisfaction with the physical education classes, using The Theory of Self-determination as a support method. The sample for this study was 922 students, of both genres and in Compulsory Secondary Education, aged between 14 and 18 years. To carry out the study, the Student’s Scale of Responsibility was used in the physical education assessment, the Basic Psychological Needs Measuring Scale, the Percentage Scale for Physical Education Causality and the Satisfaction Scale in Physical Education. The results of the structural equations model revealed a good adjustment to the data. This finding highlights the importance of giving responsibilities to the students in the evaluation process, in order to satisfy the psychological needs of the students and, therefore, self-determined motivation, thus increasing satisfaction towards physical education classes.

## Introduction

Physical education (PE) has become a framework for many youngsters to carry out physical activities, which then increases their motivation and adherence to do exercise after school [1, 2, 3].

The teacher is one of the main promotors of the practising of physical activities [4], for which their figure is crucial for the students to increase, or not, their level of regular physical activity [5]. However, in PE there are still authors [6, 7] who indicate that, traditionally, teaching has consisted in a pedagogical model of direct instruction.

Research has determined that the teacher’s use of strategies, with positive psychological aspects, such as the increase of students’ intrinsic motivation in PE classes, will allow for the development and consolidation of behaviours related to physical activity [8, 9, 10].

Motivational phenomena combining a set of biological, emotional, cognitive and social aspects, which at the same time are interrelated with each other, influence persistence, intensity and frequency of behaviour, and interact with each other by increasing, maintaining or decreasing this behaviour [11].

One of the theories that helps to explain the motivation of students in PE classes is called the Self-determination Theory (SDT) [12, 13]. The SDT, proposes that motivation is framed throughout a three-level continuum [14, 15]: autonomous motivation (the most self-determined, required for an activity to be carried out for sheer pleasure), controlled motivation (carrying out an activity for reward or recognition outside of the activity) and demotivation (the least self-determined) [16].

Furthermore, it establishes three Basic Psychological Needs (BPN): autonomy (the desire to engage in activities by one’s own choice), competence (the desire to interact efficiently with the means to feel competent) and relatedness (the desire to feel part of a group) [16, 12].

According to the SDT, the BPN constitute the psychological mediators that influence the three main types of motivation [12, 17]. Several studies have used the BPN as mediators that positively predict the more self-determined forms of motivation [18, 19, 20].

The Hierarchical Model of Motivation (HMM) [21] associates BPN with the SDT [22]. According to the HMM, the pedagogical model used by the teacher will influence the satisfaction of the BPN and, consequentially, the level of autonomous motivation of the student. The level of self-determined motivation achieved can help to positively or negatively predict the cognitive, affective and behavioural results. As a result, the students who experience positive results in PE, such as enjoyment and the intention to be physically active, present a more self-determined, and therefore more autonomous motivation [23, 24] than the students that experience negative results like boredom. It is most likely that the latter demonstrate controlled motivation or demotivation and that they will run a greater risk of giving up physical activity and sport [25]. The model establishes that the social aspects of the environment (background variables) influence motivation, depending on the achievement or not of a series of BPN (autonomy, competence and relatedness), where satisfaction increases the degree of intrinsic motivation (motivational variables) [12, 16] and will lead to positive consequences on a cognitive, affective and behavioural level (consequent variables).

In PE classes, a linear and mechanistic pedagogical model has been predominant, oriented towards the psychometric results of students at the expense of social and cognitive results [26]. However, a climate in which responsibility is given to the student will generate positive thoughts about physical activity [27]. Different studies have revealed that when the teacher provides students with autonomy and responsibility, they value more highly the PE classes and their enjoyment also increases [28, 29, 30]. In a recent study [31], with 532 students, guided by the TAD hypothesis, it was concluded that the student profiles of PE classes were mainly autonomous ones.

Evaluation can be considered as an instrument for monitoring and evaluating the results obtained by a student. The teacher can employ a more controlling teaching style, where more importance is given to results than to the learning process, or a teaching style that favours the autonomy of the student, where the student is a participant in their own learning process, using techniques such as self-evaluation, co-evaluation or hetero-evaluation [32]. Given the importance of the process in the achievement of results by the students, in this study, as a prior variable to the motivational variables, the perception of the assigning of student responsibility was used in the evaluation.

According to Hortigüela-Alcalá et al. [32], students take pleasure from being offered different strategies and alternatives to achieve their goals; This, in turn, increases intrinsic motivation towards classes and thereby the likelihood of students exercising outside of the classroom [33, 34]. Studies like that of Yonemura et al. [35], indicated that student participation in evaluation produced an increase in their commitment to learning. In the same way, it revealed the importance of proposing different strategies for evaluation in which student participation is included [36, 37]. Likewise, other authors [38, 39, 40, 41] highlight the importance of PE and sport being directed towards student autonomy and the designation of student responsibilities, claiming that a teaching style that gives subjects the chance to choose, participate and make decisions in classes, will give rise to a more enjoyable participation and an increase in intrinsic motivation [18, 42, 43]. Therefore, students need to be given the opportunity to participate, by being given responsibilities. [44, 45, 46].

Following the HMM, the consequent variable of the present study was satisfaction with PE classes. According to Herrera-Mor et al. [47], the enjoyment that is experienced from an activity, understood as satisfaction in relation to pleasure and well-being, allows for participation to remain throughout time, for a greater adherence and for participation to become an integral part of lifestyle.

Similarly, enjoyment can be understood as the valued sense of the activities carried out in PE classes by the students [48]; and this variable (satisfaction with PE classes) is even related to the obtaining of a better academic qualification [49, 50]. As some studies have indicated, to avoid the abandonment of physical activity, teachers must try to make activities fun and avoid those which are not entertaining [51], thus presenting the teacher with an essential role to play in the development of these activities [52].

In this regard, Ntoumanis [53] explained that when subjects have fun they tend to be intrinsically motivated and give more importance to the subject. Satisfaction with PE classes will be positively related to the satisfaction of BPN as well as to a more self-determined motivation [48, 52].

Moreno et al. [54], in a sample of 819 students aged between 14 and 17 years, discovered that the most self-determined form of motivation positively predicted the importance given to PE classes and with this, satisfaction with the same.

In relation to satisfaction, motivation and boredom with PE classes, as several studies have shown [10, 55, 56, 57, 58, 59], high levels of self-determination are associated with greater effort, enjoyment, the importance of PE and the development of positive behaviour. In contrast, if motivation is less self-determined, the consequences will be negative, such as boredom in classes [9, 60].

Thus, the objective for this study was to analyse the relationships between the designation of student responsibility in evaluation, motivational variations and student satisfaction with PE classes, through obtaining a model of structural equations. Specifically, the hypothesis of the study was that the perception of the assignment of student responsibility during evaluation would positively predict the satisfaction of the BPN, positively predicting autonomous motivation, which would positively predict the satisfaction with PE classes.

## Material and methods

### Research design

The study carried out was correlational of a transversal style, in which the variables described above have not been altered or manipulated, only what occurs with them under natural conditions having been observed [61].

Likewise, it is located within quantitative empirical studies and, within these, it refers to the descriptive study of populations through surveys [62].

### Sample

The study sample was 922 students of both sexes (430 male and 492 female) from compulsory secondary education, more specifically from 3rd and 4th year of secondary school.

The type of sampling that was carried out was intentional by conglomerates. Each conglomerate was constituted by a classroom of about 18-19 students, obtaining 50 conglomerates. The ages of the sample were between 14 and 18 years (M = 14.95, SD = .98).

In Table 1, the distribution of the sample in terms of gender and year-group can be seen.

**Table 1.**
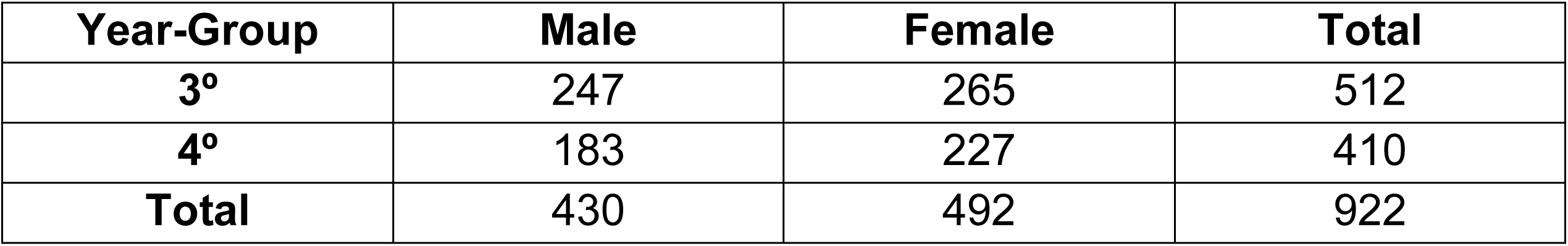
Distribution of the sample according to gender and year group.

| Year-Group | Male | Female | Total |
| --- | --- | --- | --- |
| 3° | 247 | 265 | 512 |
| 4° | 183 | 227 | 410 |
| **Total** | 430 | 492 | 922 |

In Table 2, the distribution of the sample in terms of organisations that participated in the study and year-group is represented.

**Table 2.** Distribution of the sample according to the school and the year-group.

| School | 3° |  | 4° |  | Total |
| --- | --- | --- | --- | --- | --- |
|  | Male | Female | Male | Female |  |
| School 1 | 22 | 20 | 16 | 10 | 68 |
| School 2 | 19 | 27 | 11 | 3 | 60 |
| School 3 | 22 | 36 | 24 | 32 | 114 |
| School 4 | 54 | 56 | 47 | 48 | 205 |
| School 5 | 29 | 19 | 22 | 32 | 102 |
| School 6 | 42 | 43 | 18 | 60 | 163 |
| School 7 | 18 | 21 | 0 | 0 | 39 |
| School 8 | 41 | 43 | 38 | 35 | 157 |
| School 9 | 0 | 0 | 7 | 7 | 14 |
| **Total** | 247 | 265 | 183 | 227 | 922 |
| **Total Year-Group** | 512 |  | 410 |  | 922 |

### Variables and measurement tools

In this section, the variables present in this investigation are revealed, divided according to the HMM: antecedent, motivational and consequent. In addition, a description is given of the instruments used to measure each of them.

### Antecedent variables and measurement tools

Level of responsibility of the student during evaluation: In order to know the perception of the level of responsibility that is given to the student in the evaluation, The Scale of Student Responsibility during Evaluation in Physical Education (ERAEEF) was used, adapted to Spanish by Moreno et al. [63]. It is made up of 11 items divided into 2 factors. In the present study, the factor known as the value of the transfer of responsibility in the result of the evaluation was used composed of 5 items (E.g. Working with the PE teacher to decide my score is important). Thus, in the Confirmatory Factor Analysis (CFA), the results showed acceptable adjustment indices [64]: χ^2^ = 55.00, gl = 19, p = .00, χ^2^ / df = 2.89, CFI = .99, TLI = .98, RMSEA = .05 (IC 90% = .03, .06).

### Motivational variables and measurement tools

Basic Psychological Needs: To measure the satisfaction of the BPN, the Basic Psychological Needs Measurement Scale (BPNMS) was used, the original scale of Vlachopoulos & Michailidou [65] and validated to Spanish by Moreno et al. [66]. It is composed of 12 items divided into 3 factors. Each factor is made up of 4 items: Satisfaction of the BPN of Competence (E.g. The exercises that I perform are in line with my interests), Satisfaction of the BPN of Competence (E.g. I do the exercises effectively), Satisfaction of the BPN of Relatedness (E.g. I feel that I can communicate openly with my colleagues). Regarding the CFA, the results showed acceptable adjustment indices [64]: χ^2^ = 57.79, df = 24, p = .00, χ^2^ / df = 2.41, CFI = .99, TLI = .99, RMSEA = .04 (IC 90% = .03, .05).

Levels of Self-Determined Motivation: To measure the levels of self-determined motivation, the Perceived Locus of Causality in Physical Education (PLOC) was used. Original scale by Goudas et al. [67], and validated in Spanish by Moreno et al. [13]. It consists of 20 items divided into 5 factors. In the present study, a single factor has been used, autonomous motivation, composed of the grouping of intrinsic motivation (E.g. Because I enjoy learning new skills) and identified regulation (E.g. Because it is important for me to do well in PE). Regarding the CFA, the results showed acceptable adjustment indices [64]: χ^2^ = 296.79, gl = 62, p = .00, χ^2^ / gl = 4.35, CFI = .96, TLI = .97, RMSEA = .06 (90% CI = .05, .07).

### Consequent variables and measurement tools

Satisfaction Level and Boredom in PE classes: To measure the level of satisfaction or boredom that students present in PE classes, the Basic Needs in Sport Satisfaction Scale (BNSSS) was used. A scale validated for sports by Duda & Nicholls [68] and validated to Spanish by Balaguer et al. [69]. It is composed of 8 items divided into two factors, of which only satisfaction with PE classes was used, with 5 items (E.g. Normally I find PE interesting). Thus, in the Confirmatory Factor Analysis (CFA), the results showed acceptable adjustment indices [64]: χ^2^ = 5.29, df = 2, p = .00, χ^2^ / df = 2.65, CFI = .99, TLI = .99, RMSEA = .06 (IC 90% = .04, .09).

In all of the questionnaires that were used, answers were given to all of the items through a Likert Scale of 5 points, with a range from 0, which means the student is in complete disagreement, to 5, meaning that the student completely agrees.

## Procedure

Having defined the objectives of the study, the measurement instruments were selected in order to collect information, a dossier was prepared, and some interesting data was gathered, such as age, school year, the practice of extracurricular physical activity and the school to which the students belonged. Subsequently, the different schools were contacted and the objective of the study was explained. They were given a consent form for the parents to sign, as the students were under 18 years of age.

Following this, specific days were chosen for visiting the schools and handing out the questionnaires to those subjects with parent authorisation, never in the presence of the PE teacher.

The time employed for the completion of the questionnaires was 40 minutes per class.

## Data analysis

The Confirmatory Factorial Analysis (CFA) was performed in order to verify the internal consistency of the questionnaires, and later, once the different variables were created, the descriptive statistics. The corresponding variables were created with the factors that showed adequate reliability indexes. Normality tests were performed, in order to determine what type of statistics should be used. The measurements of asymmetry, kurtosis, Kolmogorov-Smirnov, with the correction of Lilliefors, verified that the distribution of the sample was normal, for which parametric statistics were applied.

Then, Structural Equation Modelling (SEM) was employed, as it is considered to be the most effective tool for the study of causal relationships in non-experimental data [70].

The values p < .05 and p < .01 were used for statistical significance. Validity was also examined through confirmatory factor analysis, respecting the criterion of eliminating those items with a regression weight that did not present an adequate value (greater than .40) [72].

Regarding the CFA and SEM, they were carried out with Mplus [73], version 7.11. These analyses reveal some coefficients or fit indexes that allow us to check the validity of the variables of the instruments. These indexes of goodness of fit are the chi-square (χ^2^), the degrees of freedom (gl), the significance (p), the χ^2^ / gl, the RMSEA (Root Mean Square Error of Approximation), the index CFI (Comparative Fit Index) and the Tucker-Lewis Index (TLI). The χ^2^ / gl is considered acceptable when it is lower than 5, the RMSEA with values lower than .05, and the CFI and TLI with values higher than .90 [64, 74].

For the reliability, descriptive, asymmetric, kurtosis and correlation analyses, the statistical program SPSS 21.0 was used.

For the analysis of reliability, two indices were used, Cronbach’s Alpha (α) (equal to or greater than .70) [75], and Omega Coefficient (ω) [76], which also serves to check the internal consistency of the variables used in the investigation and, according to some authors (77), have shown evidence of greater accuracy. This means that in McDonald’s Omega Coefficient the established range is between 0 and 1, with the highest values giving us the most reliable measurements [77]. With the Omega Coefficient of McDonald, the calculations were made with the “psych” 1.4.2.3 [71] of R 3.0.3 (RCore-Team, 2014).

## Results

### Descriptive, reliability, asymmetry and kurtosis statistics

Table 3 presents the descriptive statistics of the instruments used for this study. This table shows the mean (M) and standard deviation (SD) of all study variables, observing that, in terms of BPN, the highest mean value was for the need for relatedness, which is equal to the satisfaction with PE classes, with the lowest average being the BPN of autonomy.

The results from the reliability analysis are also observed, in order to check the internal consistency of the questionnaires. All the factors were accepted and used in the analyses, given that Cronbach’s Alpha (α) was equal to or greater than .70 [75] and that the Omega Coefficient of McDonald has high, close to 1 [77].

**Table 3.** Reliability analysis and descriptive statistics.

| Instrument / Variable | M | TD | $\alpha$ | $\omega$ |
| --- | --- | --- | --- | --- |
| **ERAEEF** |  |  |  |  |
| Value Assignment Evaluation | 3.58 | .96 | .77 | .82 |
| **BPNES** |  |  |  |  |
| Satisfaction BPN Autonomy | 2.97 | .91 | .80 | .84 |
| Satisfaction BPN Competence | 3.68 | .95 | .80 | .84 |
| Satisfaction BPN Relatedness | 3.89 | .94 | .85 | .89 |
| **PLOC** |  |  |  |  |
| Autonomous Motivation | 3.69 | .87 | .88 | .91 |
| **SSI-EF** |  |  |  |  |
| PE Classes Satisfaction | 3.89 | .98 | .89 | .91 |

M, Media; TD, Typical Deviation; $\alpha$ , Cronbach's Alpha; $\omega$ , Omega coefficient.

In accordance with the normality rules proposed by Curran et al. [78] all of the variables comply with the univariate normality, since the values of asymmetry were below 2 and those of kurtosis below 7. These results can be seen in Table 4.

**Table 4.**
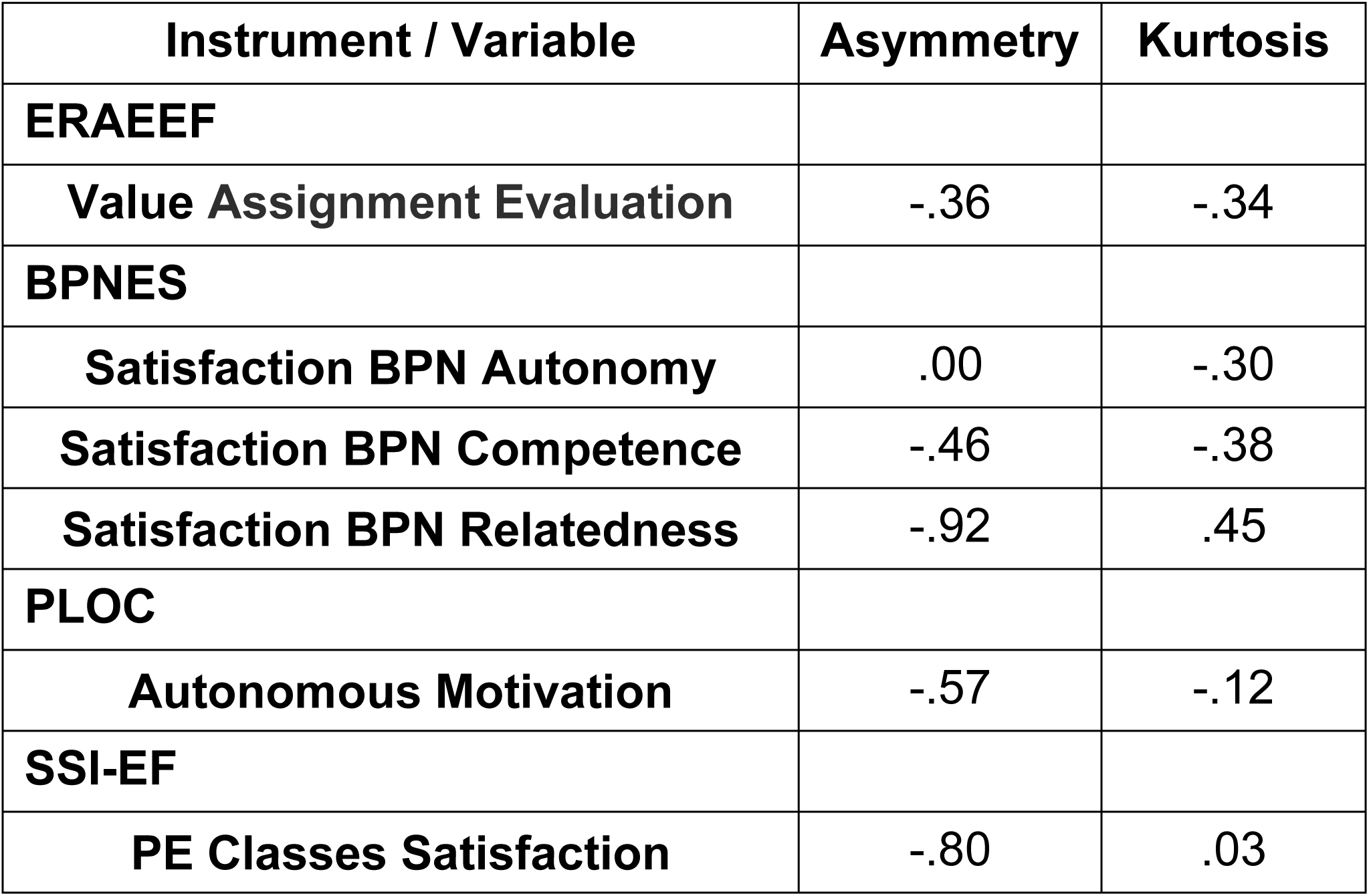
Asymmetry and kurtosis of the variables under study.

| **Instrument / Variable** | **Asymmetry** | **Kurtosis** |
| --- | --- | --- |
| **ERAEEF** |  |  |
| **Value Assignment Evaluation** | -.36 | -.34 |
| **BPNEs** |  |  |
| **Satisfaction BPN Autonomy** | .00 | -.30 |
| **Satisfaction BPN Competence** | -.46 | -.38 |
| **Satisfaction BPN Relatedness** | -.92 | .45 |
| **PLOC** |  |  |
| **Autonomous Motivation** | -.57 | -.12 |
| **SSI-EF** |  |  |
| **PE Classes Satisfaction** | -.80 | .03 |

### Analysis of structural equations

In line with the HMM [21, 79], the antecedent variables (perception of the transfer of responsibility to the student in evaluation), the mediators (satisfaction of the BPN), self-determined types of motivation (autonomous motivation, controlled motivation and demotivation) and consequences (satisfaction with PE classes) were included.

In this model, the aim was to find out the predictors of satisfaction with PE classes, based on the perception of assigning responsibility to students and the motivational variables (satisfaction of BPN and autonomous motivation). The results are shown in Fig. 1.

**Fig. 1.**
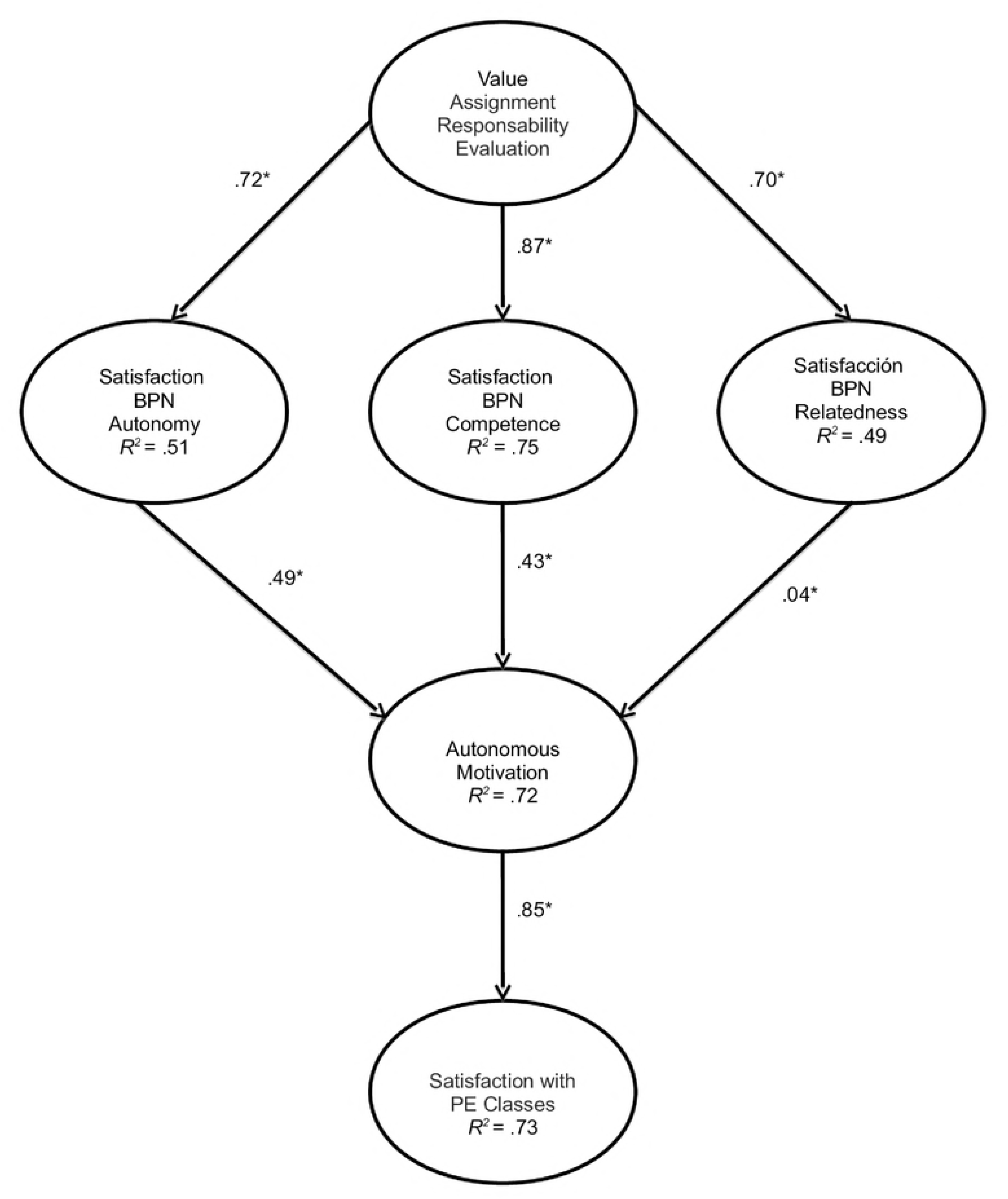
SEM. Predicting the students’ satisfaction with PE classes from assigning them with responsibility in evaluation and motivational variables. All of the parameters are standardised, the most statistically significant are indicated with *p < .01.

The contribution of each of the factors to the prediction of other variables was examined through the standardised regression weights, hence, the value of the assignment of responsibility in the result of the evaluation predicted in a positive and significant way the satisfaction of the BPN of autonomy (β = .72), competence (β = .87) and relatedness (β = .70). On the other hand, autonomous motivation was predicted in a positive way by the satisfaction of the BPN of autonomy (β = .49), competence (β = .43) and relatedness (β = .04), with self-motivation predicting in a positive and meaningful way the satisfaction with PE classes (β = .85).

The results of the model of structural equations revealed a good alignment to the data [64, 74], as can be seen in Table 5.

**Table 5.** Adjustment indices of the Structural Equation Modelling.

| Indices | Values |
| --- | --- |
| $\chi^2$ | 1004.16 |
| gl | 202 |
| p | .00 |
| $\chi^2/\text{gl}$ | 4.93 |
| CFI | .94 |
| TLI | .93 |
| RMSEA | .06 (IC 90% = .06, .07) |

Regarding the indirect effects between the latent variables, the results are shown in the Table 6.

**Table 6.** Indirect effects in Structural Equation Model.

| Variables | Effects |
| --- | --- |
| Value Assignment Responsibility Evaluation → Autonomous Motivation | .75 |
| Value Assignment Responsibility Evaluation → Classes Satisfaction | .64 |
| Satisfaction BPN Autonomy → Classes Satisfaction | .42 |
| Satisfaction BPN Competence → Classes Satisfaction | .37 |
| Satisfaction BPN Relatedness → Classes Satisfaction | .03 |

Table 6 shows the totals of the indirect effects. However, the results demonstrate that the indirect effects of the assigning of student responsibility in evaluation upon autonomous motivation vary according to the production level, via the satisfaction of the BPN of autonomy, which was β=.35. The satisfaction of the BPN of competence being β=.37 and the satisfaction of the BPN of relatedness β=.03. Regarding the variable for the value of assigning students with responsibility during evaluation, we obtained that, via the satisfaction of students in PE classes and autonomous motivation, β=.30, via the satisfaction of the BPN of competence β=.32 and, via the satisfaction of the BPN of relations and autonomous motivation β=.02.

## Discussion and conclusions

Given the relevance of assigning responsibility to the student in the evaluation for improving motivational processes and increasing student satisfaction with PE classes, the present study aimed to validate a model that would analyse these relationships from the HMM. The hypothesis stated that the perception of the assignment of responsibility to the student during evaluation would positively predict the satisfaction of the BPN, which would positively predict autonomous motivation, in turn positively predicting the satisfaction with PE classes.

In terms of the results previously mentioned, It can be observed that the hypothesis was fulfilled, as it was already demonstrated that the perception of the assignment of responsibility to the student in the evaluation predicted in a positive and significant way the satisfaction of the BPN (autonomy, competence and relatedness), and that the satisfaction of the BPN positively predicted autonomous motivation, with a significant prediction of the BPN of autonomy and competence. Finally, the most self-determined form of motivation positively and significantly predicted satisfaction with PE classes.

In line with the model obtained, Moreno et al. [80] showed that responsibility positively predicted psychological mediators, and this predicted intrinsic motivation, which positively predicted the importance that students give to physical education, and this, finally, positively predicted the student’s intention to continue playing sports.

Other studies have demonstrated that when students are offered the opportunity to choose tasks they improve their skills, their physical activity and their perceived competence [81]. It was also proven that there was a greater learner involvement when given the opportunity to make decisions with various methodological aspects such as space, time, material or grouping [82].

Motivation involves a set of emotional, cognitive and social phenomena, with which, according to studies, if a teaching style is used where students are allowed to participate in the teaching-learning process, the cognitive and physical involvement will be greater [83]. This explains a greater satisfaction towards PE classes, and a greater commitment to learning, as students are more intrinsically motivated thanks to their involvement in the evaluation process [35]. Research carried out by Vera [30], with 49 students, also showed that the assignment of responsibilities to the students makes the satisfaction of the BPN of autonomy higher, and with it the satisfaction and enjoyment towards physical activity.

It has been confirmed that one key aspect to improving motivation is the assignment of responsibility to the student [18], along with the use of styles that favour the autonomy of the students [84]. Thanks to different works [18, 42], which are in line with our results, it can be affirmed that an assignment of responsibilities increases the most self-determined forms of motivation. The study by Gómez-Rijo et al. [85], reached the conclusion that the transfer of responsibilities to the student, by the teacher, contributes to the development of student autonomy. It also demonstrated [85] that giving autonomy to the student for the learning of physical skills improves autonomous motivation.

Evaluation must not only be linked to the teacher giving a score, but the student must be given the possibility to decide and intervene, taking into account initial and bidirectional agreements [87]. Other works developed in the educational field [88, 89], that related the satisfaction of the BPN to the self-determined forms of motivation, revealed, like in our study, that an adequate satisfaction of the BPN would increase intrinsic motivation.

The authors of these works have shown that a greater feeling of autonomy will increase intrinsic motivation [8, 9, 53, 60, 90, 91] and with this, satisfaction when participating in activities [39, 40, 41], supporting the results found in this study. However, other authors, performing intervention programmes to support teachers with the BNP, did not find significant results in intrinsic motivation [91].

A teaching-style where autonomy and decision-making is stimulated will reduce the demotivation of the students, as well as boredom with PE classes [94], as pointed out in the study by Moreno et al. [93]. Different works have indicated that the less self-determined forms of motivation and a lower perception of satisfaction of BPN [95] are related to the giving up of physical activity, which may be due to the lack of satisfaction with the PE classes.

Other studies which had similar results to ours, indicate that an increase in autonomy will make satisfaction with PE classes higher [82], and that students will be more involved in their tasks and their own learning process [82]. Different investigations, such as the one carried out by Méndez et al. [94], found that if a suitable atmosphere that involves the task is generated in the classroom, the satisfaction of the BPN will be greater, which will be positively related to more self-determined motivation and with less boredom with the classes of PE.

Research that is also related to our variables [48, 52, 54, 55], indicated that the satisfaction of the BPN predicted high levels of intrinsic motivation and that this was related to an increase in enjoyment and satisfaction with classes. It can therefore be said that there is a close relationship between intrinsic motivation and satisfaction with classes [24, 96]. Some authors, in their results give particular importance to the BPN of autonomy [61, 98].

As many researchers have been proposing for some time [36, 37, 98], it is necessary to come up with new evaluation strategies which offer more student involvement. Strategies such as developing the students’ ability to reflect on what they have done, substitute the final exam for a continuous process in which the students learn from their mistakes and successes, involve the student in making decisions, among others, which will mean that, based on the theoretical postulates of the HMM, the satisfaction of the BPN will be higher, as well as that the more self-determined forms of motivation will be increased, with levels of demotivation decreasing. This will have positive consequences, such as satisfaction with PE classes, and therefore, increase the possibility of physical activity outside the classroom.

One of the limitations found in this study was the sample, which would be interesting to expand to other areas and even differentiate by age, gender and socioeconomic level. Another limitation was seen from only using questionnaires, as only opinion is determined through a scale of answers. It would be interesting to make a methodological triangulation, using systematic observation and the use of interviews, both with students and with teachers. Once the results are known, a longitudinal or quasi-experimental study could be carried out, through an intervention that would allow us to establish cause-effect relationships, in order to know the effect caused by the application of different motivational strategies in the variables under analysis.

In conclusion, thanks to results from models such as ours, in PE classes intervention programmes are necessary to achieve more self-determined motivation of our students, using different strategies such as posing tasks that are fun for them, by assigning them with responsibilities, proposing self-evaluation activities, as well as reciprocal evaluation so that they feel they are participants in their teaching and learning process. In this way, we can achieve that the students increase their levels of satisfaction of the BPN, which will lead to them showing higher levels of autonomous motivation in the classes and, consequently, to their satisfaction with the PE classes being greater.

Author Contributions
Conceived and designed the experiments: ML SL RJ. Performed the experiments: ML SL MB RJ. Analysed the data: SL RJ. Contributed analysis tools: ML SL MB RJ. Wrote the paper: ML.

## References

1. Franco E, Coterón J, Martínez HA, Brito J. Motivational profiles in physical education students from three countries and their relationship with physical activity. Suma Psicol. 2017; 24: 1–8. http://dx.doi.org/10.1016/j.sumpsi.2016.07.001

2. Pulido JJ, Sánchez-Oliva D, Amado D, González-Ponce I, Sánchez-Miguel PA. Influence of motivational processes on enjoyment, boredom and intention to persist in young sportspersons. S Afr J Res Sport PH. 2014; 36(3): 135–49.

3. Taylor IM, Spray C, Pearson N. The influence of thephysical education environment on children’s well-being and physical activity across the transition from primary to secondary school. J Sport Exerc Psychol. 2014; 36: 574–83. http://dx.doi.org/10.1123/jsep.2014-0038.

4. Egan CA, Webster CA. Using Theory to Support Classroom Teachers as Physical *Activity Promoters*. J Physical Educ, Recre Dance. 2018; 89(1): 23–9.

5. Yilmaz A, Esenturk OK, Demir GT, Ilhan EL. Metaphoric Perception of Gifted Students about Physical Education Course and Physical Education Teachers. J Educ Learn. 2017; 6(2): 220–34.

6. Gil-Arias A, Harvey S, Cárceles A, Práxedes A, Del Villar F. Impact of a hybrid TGfU-Sport Education unit on student motivation in physical education. PloS ONE. 2017; 12(6): e0179876.

7. Ayuso JAZ. Benefits of teaching styles and student-centred methodologies in Physical Education. E-Balonmano. 2018; 13(3): 237–50.

8. Sánchez-Oliva D, Pulido-González JJ, Leo FM, González-Ponce I, García-Calvo T. Effects of an intervention with teachers in the physical education context: A Self-Determination Theory approach. PloS ONE. 2017; 12(12): e0189986.

9. Standage M, Duda JL, Ntoumanis N. A test of Self-Determination Theory in school physical education. Brit J Educ Psychol. 2005; 75: 411–33. pmid:16238874.

10. Taylor IM, Ntoumanis N, Standage M, Spray CM. Motivational predictors of physical education students’ effort, exercise intentions, and leisure-time physical activity: A multilevel linear growth analysis. J Sport Exerc Psychol. 2010; 32(1): 99–120.

11. Escartí A, Cervelló EM. Motivation in Sport. In I. Balaguer (Ed.), Psychological training in sport: Principles and applications (pp. 61–90). Valencia: Albatros Educación; 1994.

12. Ryan RM, Deci EL. Self-determination theory and the facilitation of intrinsic motivation, social development, and well-being. Am Psychol. 2000; 55: 68–8. pmid:11392867

13. Moreno JA, González-Cutre D, Chillón M. Preliminary validation in Spanish of a scale designed to measure motivation in physical education classes: the Perceived Locus of Causality (PLOC) Scale. Spa J Psychol. 2009; 12: 327–37.

14. Vansteenkiste M, Lens W, Deci EL. Intrinsic versus extrinsic goal contents in self-determination theory: Another look at the quality of academic motivation. Educ Psychol. 2006; 41(1): 19–31. doi:10.1207/s15326985ep4101_4

15. Vansteenkiste M, Niemiec C, Soenens B. The development of the five mini-theories of self-determination theory: An historical overview, emerging trends, and future directions. In T. Urdan & S. Karabenick (Eds.). Advances in Motivation and Achievement, vol. 16: The decade ahead (pp.105–166). Bingley, UK: Emerald; 2010. doi:10.1108/S0749-7423(2010)000016A007.

16. Deci EL, Ryan RM. The ‘‘what” and ‘‘why” of goal pursuits: human needs and the self-determination of behaviour. Psychol Inq. 2000; 11: 227–68.

17. Deci EL, Ryan RM. Intrinsic motivation and Self-determination in human behavior. New York: Plenum; 1985.

18. González-Cutre D, Sicilia A, Moreno JA. A Quasi-experimental Study of the Effects of Task-involving Motivational Climate in Physical Education Classes. Rev Educ. 2011; 356: 677–700.

19. McDonough MH, Crocker PRE. Testing self-determined motivation as a mediator of the relationship between psychological needs and affective and behavioral outcomes. J Sport Exerc Psychol. 2007; 29(5): 645–63.

20. Standage M, Duda JL, Ntoumanis N. Students motivational processes and their relationship to teacher ratings in school physical education: A self-determination theory approach. Res Q Exerc Sport. 2006; 77(1): 100–10.

21. Vallerand RJ. A hierarchical model of intrinsic and extrinsic motivation in sport and exercise. In G. Roberts (Ed.). Advances in motivation in sport and exercise (2nd ed., pp.263–319). Champaign, IL: Human Kinetics; 2001.

22. Deci EL, Ryan RM. Handbook of self-determination research. Rochester, New York: University of Rochester Press; 2002.

23. Gucciardi DF, Jackson B. Understanding sport continuation: An integration of the theories of planned behaviour and basic psychological needs. J Sci Med Sport. 2015; 18(1): 31–6. pmid:24378719.

24. Sánchez-Oliva D, Sánchez-Miguel PA, Leo FM, Kinnafick FE, García-Calvo T. Physical education lessons and physical activity intentions within Spanish secondary schools: a self-determination perspective. J Teach Phys Educ. 2014; 33: 232–49.

25. Ntoumanis N, Standage M. Motivation in physical education classes: A self-determination theory perspective. Theory Res Educ. 2009; 7: 194–202.

26. Light R, Fawns R. Knowing the game: Integrating speech and action in games teaching through TGfU. Quest. 2003; 55(2): 161–76.

27. Derry JA. Single-sex and coeducation physical education: perspective of adolescent girls and female physical education teachers (research). Melpomene Journal. 2002; 22: 17–28.

28. Allen JB. Social motivation in youth sport. J Sport Exerc Psychol. 2003; 25: 551–67.

29. Guan J, Xiang P, McBride R, Bruene A. Achievement goals, social goals and students’ reported persistence and effort in high school physical education. J Teach Phys Educ. 2006; 25: 58–74.

30. Vera JA. Dilemmas in the negotiation of curriculum to students from the trasnfer of responsability for the evaluation in Physical Education classroom. Rev Inv Educ. 2010; 7: 72–82.

31. Bechter BE, Dimmock JA, Howard JL, Whipp PR, Jackson B. Student motivation in high school physical education: A latent profile analysis approach. J Sport Exerc Psychol. 2018; 40(4): 206–16.

32. Hortigüela-Alcalá D, Pérez-Pueyo Á, Fernández-Río J. Connection between the attitudinal style and students’ assessment responsibility in Physical Education. Cul Cien Dep. 2017; 12(35): 89–99

33. Baena A, Gómez M, Granero A, Ortíz MM. Predicting satisfaction in physical education from motivational climate and self-determined motivation. J Teach Phys Educ. 2015; 34(2): 210–24. doi.org/10.1123/jtpe.2013-0165.

34. Hassandra M, Goudas M, Chroni S. Examining factors associated with intrinsic motivation in physical education: a qualitative approach. Psychol of Sport Exerc. 2003; 4: 211–23.

35. Yonemura K, Fukugusakio Y, Yoshinaga T, Takahashi T. Effects of Momentum and climate in Physical Education class on students´ formative evaluation. Int J Sport Health Sci. 2003; 2: 25–33.

36. Álvarez-Méndez JM. The formative evaluation. Cuad Pedag. 2007; 364: 96–100.

37. Vera JA, Moreno JA. The teaching of responsibility in the physical education classroom. Habilidad Motriz. 2008; 32: 39–43.

38. Moreno JA, Vera JA, Del Villar F. Search for autonomy in motor task learning in physical education university students. Eur J Psychol Educ. 2010; 25(1): 37–47.

39. Prusak KA, Treasure DC, Darst PW, Pangrazi RP. The effects of choice on the motivation of adolescent girls in physical education. J Teach Phys Educ. 2004; 23: 19–29.

40. Wallhead TL, Ntoumanis N. Effects of a sport education intervention on students’ motivational responses in physical education. J Teach Phys Educ. 2004; 23: 4–18.

41. Ward P. What we teach is as important as how we teach it. J Physical Educ, Recre Dance. 2006; 77: 20–3.

42. Moreno JA, Sicilia A, Sáenz-López P, González-Cutre D, Almagro BJ, Conde C. Motivational analysis comparing three contexts of physical activity. Rev Int Med Cienc Ac. 2014; 14(56): 665–85.

43. Moreno JM, Vera JA. Model Causal of the Satisfaction with the Life in Adolescent Students of Physical Education. Rev Psic. 2011; 16(2): 367–80.

44. Hellison D. Teaching personal and social responsibility in physical education. En S. J. Silverman y C. D. Ennis (Eds.), Student learning in physical education: applying research to enhance instruction (pp. 241–254). Champiagn: Human Kinetics; 2003.

45. Hellison D, Martinek T. Social and individual responsibility programs. En D. Kirk, D. Mcdonald & M. O´Sullivan (Eds), The handbook of physical education (pp. 610–626). Thousand Oaks, CA: Sage; 2006.

46. Ruiz LM, Rodríguez P, Martinek T, Schilling T, Durán LJ, Jiménez P. Project Effort: A model for the Development of Social and Personal Responsibility through Sport. Rev de Educ. 2006; 341: 933–58.

47. Herrera-Mor E, Pablos-Monzó A, Chiva-Bartoll O, Pablos-Abella C. Effects of a global physical activity program on the physical condition, self-esteem and enjoyment on elderly adults. Ágora. 2016; 18(2): 167–83.

48. Moreno JA, González-Cutre D, Ruiz LM. Selfdetermined motivation and physical education importance. Hum Mov. 2009; 10(1): 5–11.

49. Moreno JA, Silveira Y, Alias A. Predictive model to improve the competence perception and academic performance in colleges. REDU. Rev Doc Univ. 2015; 13: 173–88.

50. Sevil J, Aibar A, Abós A, García L. Motivational climate of teaching physical education: Could it affect student grades?. Retos. 2017; 31: 98–102.

51. Ortega E, Calderón A, Palao JM, Puigcerver C. Design and validation of a questionnaire to evaluate the perceived attitude of the professor and of a questionnaire to evaluate the actitudinal contents of the students during the physical education classes in secondary education. Retos. 2008; 14: 22–9.

52. Sevil J, Abós A, Generelo E, Aibar A, García-González L. Importance of support of the basic psychological needs in predisposition to different contents in Physical Education. Retos. 2016; 29: 3–8.

53. Ntoumanis N. A prospective study of participation in optional school physical education using a Self-Determination Theory framework. J Educ Psychol. 2005; 97: 444–53. doi:10.1037/0022-0663.97.3.444

54. Moreno JA, Zomeño T, Marín LM, Ruiz LM, Cervelló E. Perception of the usefulness and importance of physical education according to motivation generated by the teacher. Rev Educ. 2013; 362: 380–401.

55. Grasten A, Jaakkola T, Liukkonen J, Watt, A, Yli-Piipari S. Prediction of the enjoyment in school physical education. J Sport Sci Med. 2012; 11: 260–69.

56. Cheon SH, Reeve J, Ntoumanis N. A needs-supportive intervention to help PE teachers enhance students’ prosocial behavior and diminish antisocial behavior. Psychol Sport Exerc. 2018; 35: 74–88.

57. Moy B, Renshaw I, Davids K. The impact of nonlinear pedagogy on physical education teacher education students’ intrinsic motivation. Phys Educ Sport Pedagog. 2016; 21(5): 517–38.

58. Sánchez-Oliva D, Leo FM, Sánchez-Miguel PA, Amado D, García-Calvo T. Development of a causal model to explain positive behaviors in physical education classes. Acción Motriz. 2013; 10: 48–58.

59. Granero-Gallegos A, Baena-Extremera A, Sánchez-Fuentes JA, Martínez-Molina M. Motivational profiles of autonomy support, self-determination, satisfaction, importance of physical education and intention to partake in leisure time physical activity. Cuad Psicol Deporte. 2014; 14(2): 59–70.

60. Ntoumanis N. A self-determination approach to the understanding of motivation in physical education. Br J Educ Psychol. 2001; 71: 225–42.

61. Cubo S, Martín B, García JL. Research methods and data analysis in social sciences and health. Madrid: Ediciones Pirámide Grupo Anaya, S.A.; 2011.

62. Montero I, León OG. A guide for naming research studies in Psychology. INT J Clin Health Psychol. 2007; 7(3): 847–62.

63. Moreno JA, Vera JA, Cervelló E. Participative evaluation and responsibility in physical education. Rev Educ. 2006; 340: 731–54.

64. Hu L, Bentler PM. Cutoff criteria for fit indexes in covariance structure analysis: Conventional criteria versus new alternatives. Struct Equ Model. 1999; 6: 1–55.

65. Vlachopoulos SP, Michailidou S. Development and initial validation of a measure of autonomy, competence, and relatedness in exercise: The Basic psychological needs in exercise scale. Meas Phys Educ Exerc Sci. 2006; 103: 179–201.

66. Moreno JA, González-Cutre D, Chillón M, Parra N. Adaptation of the Basic Psychological Needs in Exercise Scale to Physical Education. Rev Mex Psicol. 2008; 25: 295–303.

67. Goudas M, Biddle SJH, Fox KR. Achievement goal orientations and intrinsic motivation in physical fitness testing with children. Pediatr Exerc Sci. 1994; 6; 159–67.

68. Duda JL, Nicholls JG. Dimensions of achievement motivation in schoolwork and sport. J Educ Psychol. 1992; 84(3): 290–99.

69. Balaguer I, Atienza FL, Castillo I, Moreno Y, Duda JL. Factorial structure of measures of satisfaction/interest in sport and classroom in the case of Spanish adolescents. Abstracts of 4th European Conference of Psychological Assessment (p. 76). Lisbon: Portugal; 1997.

70. Aron A, Aron E. Statistics for Psychology. Buenos Aires: Pearson Education; 2001.

71. Mullan E, Markland D, Ingledew DK. A graded conceptualisation of selfdetermination in the regulation of exercise behaviour: Development of a measure using confirmatory factor analytic procedures. Personal Individ Differ. 1997; 23: 745–52. doi:10.1016/S0191-8869(97)00107-4.

72. Revelle W. Psych: Procedures for Psychological, Psychometric, and Personality Research. Illinois: Evanston; 2014. Retrieved from http://cran.r-project.org/package=psych

73. Muthén LK, Muthén BO. Mplus User’s Guide (7th ed.). Los Angeles, CA: Muthén & Muthén; 2014.

74. Kline RB. Principles and practice of structural equation modeling. Structural Equation Modeling. New York: Guilford Press; 2011. http://doi.org/10.1038/156278a0.

75. Nunnally JC. Psychometric theory. New York: McGraw-Hill; 1978.

76. McDonald RP. Test theory. A unified treatment. Mahwah, NJ, Lawrence Erlbaum Associates; 1999.

77. Revelle W, Zinbarg RE. Coefficients Alpha, Omega, and the Gbl: Comments on Sijtsma. Psychometrika. 2009; 74(1): 145–54. http://doi.org/10.1007/s11336–008–9102-z

78. Curran PJ, West SG, Finch JF. The robustness of test statistics to nonnormality and specification error in Confirmatory Factor Analysis. Psychol Methods. 1996; 1: 16–29.

79. Vallerand RJ. Toward a hierarchical model of intrinsic and extrinsic motivation. En M. P. Zanna (Ed.), Advances in experimental social psychology (pp.271–360). Academic Press: New York; 1997.

80. Moreno JA, Huéscar E, Cervelló E. Prediction of adolescents doing physical activity after completing secondary education. Spa J Psychol. 2012; 15(1): 90–100.

81. Hastie PA, Rudisill ME, Wadsworth DD. Providing students with voice and choice: lessons from intervention research on autonomy-supportive climates in physical education. Sport Educ and Soc. 2013; 18: 38–56.

82. Calderón A, Martínez D, Hastie P. Students and teachers’ perception after practice with two pedagogical models in Physical Education. RICYDE. 2013; 9(32): 137–53. http://dx.doi.org/10.5232/ricyde2013.03204

83. Sánchez B, Byra M, Wallhead T. Students’ perceptions of the command, practice, and inclusion styles of teaching. Phys Educ Sport Pedagog. 2012; 17(3): 317–30. http://dx.doi.org/10.1080/17408989.2012.690864

84. Reeve J, Vansteenkiste M, Assor A, Ahmad I, Cheon SH, Jang H, Kaplan H, Moss J, Olaussen B, Wang CKJ. The beliefs that underlie autonomy supportive and controlling teaching: A multinational investigation. Motiv Emot. 2014; 38(1): 93–110. http://dx.doi.org/10.1007/s11031-013-9367-0.

85. Gómez-Rijo A, Jiménez-Jiménez F, Sánchez-López CR. Development of Autonomy of Elementary Students in Physical Education through a process of action research. RICYDE. 2015; 42(11): 310–28. http://dx.doi.org/10.5232/ricyde2015.04201

86. Behzadnia B, Mohammadzadeh H, Ahmadi M. Autonomy-supportive behaviors promote autonomous motivation, knowledge structures, motor skills learning and performance in physical education. Curr Psychol. 2017; 1–14. https://doi.org/10.1007/s12144-017-9727-0.

87. Papinczak T, Babri A, Peterson R, Kippers V, Wilkinson D. Students Generating Questions for Their Own Written Examinations. Adv Health Sci Educ. 2011; 16(5): 703–10.

88. Méndez JI, Fernández-Río J. Social responsibility, basic psychological needs, intrinsic motivation, and friendship goals in physical education. Retos. 2017; 32: 134–39.

89. Núñez JL, León J. The Mediating Effect of Intrinsic Motivation to Lear non the Relationship between Student’s Autonomy Support and Viality and Deep Learning. Spa J Psychol. 2016; 19(42): 1–6. http://doi.org/10.1017/sip.2016.43

90. Taylor IM, Ntoumanis N. Teacher motivational strategies and student self-determination in physical education. J Educ Psychol. 2007; 99(2): 747–60. doi:10.1037/0022-0663.99.4.747

91. Amado D, Del Villar F, Leo FM, Sánchez-Oliva D, Sánchez-Miguel PA, García-Calvo T. Effect of a multi-dimensional intervention programme on the motivation of physical education students. PLoS ONE. 2014; 9(1): e85275.

92. Tessier D, Sarrazin P, Ntoumanis N. The effect of an intervention to improve newly qualified teachers’ interpersonal style, students motivation and psychological need satisfaction in sport-based physical education. Contemp Educ Psychol. 2010; 35: 242–53

93. Moreno JA, Parra N, González-Cutre D. Influence of autonomy support, social goals and relatedness on amotivation in physical education classes. Psicoth. 2008; 20(4): 636–41.

94. Méndez A, Fernández J, Cecchini JA. Motivational climates, needs, motivation and outcomes in Physical Education. Aula Abierta. 2013; 41(1): 63–72.

95. Cervelló E. Psychological variables related to the choice of sports tasks with different level of difficulty. Considerations for the design of motivational programs of psychological training in sport. Eur J Hum Mov. 1999; 5: 35–52.

96. Sparks C, Lonsdale C, Dimmock J, Jackson B. An intervention to improve teachers’ interpersonally involving instructional practices in high school physical education: Implications for student relatedness support and in-class experiences. J Sport Exerc Psychol. 2017; 39(2): 120–33.

97. Baena A, Gómez-López M, Granero-Gallegos A, Martínez-Molina M. Prediction model of satisfaction and enjoyment in Physical Education from the autonomy and motivational climate. Universitas Psychologica. 2016; 15(2): 15–25. doi: 10.11144/Javeriana.upsy15-2.mpsd

98. Lewis R. Classroom discipline student responsibility: the students’ view. Teach Teacher Educ. 2001; 17: 307–19.

